# High dimensional tissue specific regulatory codes embedded in viral coding regions

**DOI:** 10.64898/2025.12.27.696677

**Authors:** Alma Davidson, Marina Parr, Franziska Totzeck, Alexander Churkin, Danny Barash, Dmitrij Frishman, Tamir Tuller

## Abstract

Viruses co-evolve with their hosts and adapt various genomic level strategies to ensure their fitness. Previous studies have demonstrated that viruses share high dimensional ‘silent’ patterns with their hosts.

In this study, based on a novel computational measure, we demonstrated for the first time that this high dimensional information encodes regulatory codes related to viral tropism.

## Introduction

Viruses depend on host-cell machinery for particle assembly ^1,2^, genome replication ^2,3^, and gene expression ^4^ to achieve efficient infection and spread. They have co-evolved with their hosts over millions of years, both to escape host immune system and to establish optimal conditions for viral fitness ^5,6^. Viral evolution is rapid because viruses have short generation times and large population sizes that lead to high mutation rates ^7,8^. Thus, viruses can rapidly generate *de-novo* mutations, producing large genetic variant pools from which selection favors more adaptive ones. Natural selection favors adaptive mutations that enhance viral replication and increase protein-synthesis efficiency ^7–10^. Frequent virus-host interactions have led to convergence in codon-usage patterns that are preserved, optimized, and traceable across viral evolution ^11–13^.

Codon usage bias (CUB) is a common feature of viruses and serves as an important indicator of their adaptation to their host ^14–20^. A study conducted a comprehensive analysis of human RNA viruses, comparing their codon usage bias to expected codon frequencies based on host usage. The results revealed a correlation between codon bias and viral structure in aerosol-transmitted viruses, suggesting the presence of translational selection within these viruses. Another study examined the impact of dinucleotide composition on codon usage bias and its implications for translational efficiency, providing additional evidence for host-driven selection ^21,22^.

Viruses have evolved over time to adapt to their host’s cellular environment. However, in multicellular organisms such as humans, the cellular environment varies among different tissues ^23,24^. Therefore, viruses have developed the ability to target specific cell types within particular tissues, which is defined as tissue tropism ^25^. This adaptability may be restricted to a single tissue in some viruses, while others are capable of infecting multiple tissues ^26,27^. For example, the human immunodeficiency virus (HIV) primarily targets immune system cells such as T cells and macrophages ^28^, while severe acute respiratory syndrome coronavirus 2 (SARS-CoV-2) has been reported to infect multiple organs in the human body, such lung ^29,30^, trachea ^30^, small intestine ^29^, kidney ^29–31^, pancreas ^29,31^, heart ^29,30^, adrenal gland ^30,31^, testicles ^30,31^, ovaries ^30^ and nervous system ^29,31^.

The ability to infect a specific tissue depends on various factors, such as the compatibility between viral proteins and host cell receptors ^32–34^, differences in tissue microenvironments ^35^, and the presence of transcription factors required for viral replication ^36,37^. Furthermore, differences in gene expression between tissues can influence viral infection and its ability to complete its replication cycle ^38^.

The diverse biological characteristics of human viruses may also be reflected in their codon usage bias (CUB). Such variation in CUB may be driven by differences in tRNA availability and translational efficiency across tissues ^39–41^. However, standard CUB metrics model only the independent frequencies of individual codons. High dimensional nucleotide patterns, such as substrings longer than single codons, form the basis of many other features that can capture aspects of gene expression regulation ^42,43^. Combining codon-level and high-dimensional features can provide a more comprehensive analysis of translational and regulatory dynamics.

In this study, we aim to identify patterns in the genomes of human-infecting viruses that may be indicative of their tissue tropism. While previous studies on this topic have focused on shorter sequence features such as codon usage bias and CpG content, we aim to explore nucleotide patterns longer than a codon. Our goal is to utilize these patterns, which may be involved in viral gene regulation and tissue-specific adaptation, to help predict viral tissue tropism.

## Methods

### Data

#### Human data extraction

Human transcript data were obtained from Human Genome Resources at NCBI (RefSeq Transcripts, GRCh38 edition) ^44^. Codon Usage tables were obtained from Codon and Codon Pair Usage Tables (CoCoPUTs) ^45^.

#### Human per-tissue gene expression

Isoform expression levels in human genes were obtained from The Cancer Genome Atlas (TCGA) database (https://www.cancer.gov/tcga). The publicly available gene expression quantification was filtered to include only healthy samples across 23 different tissues: BLCA, BRCA, CESC, CHOL, COAD, ESCA, GBM, HNSC, KICH, KIRC, KIRP, LIHC, LUAD, LUSC, PAAD, PCPG, PRAD, READ, SARC, SKCM, STAD, THCA, THYM and UCEC.

Because each tissue includes multiple patient samples, the median transcript per million (TPM) levels across samples was used to represent the expression level in each tissue.

Tissue-specific CU was calculated as a weighted sum of each codon counts and their TPM values from TCGA:

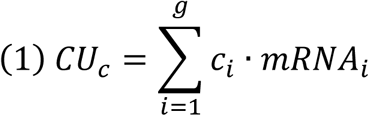

where g is the number of genes analyzed, c_i_ is the count of codon c in gene i, and mRNA_i_ is the TPM levels of gene i.

#### tRNA copy numbers and s-values

tRNA copy number, used as an estimation for the number of available tRNA molecules, was obtained from GtRNAdb database for Homo sapiens (GRCh38/hg38) edition, which includes tRNA gene predictions generated by tRNAscan-SE ^46,47^. Tissue specific tRNA levels were retrieved from the tRNA abundances quantified in ^48^.

s-values are selective constraints on the efficiency of the codon–anticodon coupling, which can vary and be optimized among different organisms. Human specific s-values were obtained from the optimized values in ^49^.

#### Tropism-defined human viruses

Data for 987 human virus species, comprising 11,573 viral proteins, were obtained from the ViralZone database ^50^. The dataset included Baltimore classification, family, and tropism annotations where available. 445 species with an available tropism were grouped into six categories based on the closest matching cell on ViralZone: Liver, Gastrointestinal tract, Immune system, Mucosal epithelium, Nervous system, and Respiratory tract.

### Codon Adaptation Metrics

#### Codon Adaptation Index (CAI)

The index measures the codon usage within highly expressed genes in a reference genome and compares the frequency of each codon to that of its synonymous alternatives. By assigning a specific weight to each codon in the human genome and then calculating the total score for viral genes, we estimated the degree of adaptation of those genes to their human hosts ^51^.

Relative synonymous codon usage for each codon within the human genome was calculated as follows:

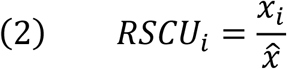

where *x*_*i*_ is the observed frequency of codon i which encodes to amino acid x, and *x̂* is the expected frequency of the codon, both in the human genome. The expected frequency is calculated:

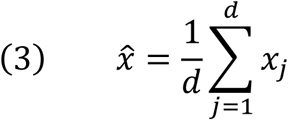

where d is the number of types of codons that encode to x amino acids. Then, the codon adaptation index for codon *i* is calculated relatively to the most frequent synonymous codon:

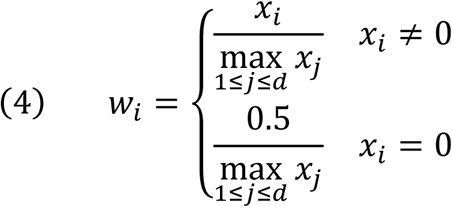

#### tRNA Adaptation Index (tAI)

The tRNA Adaptation Index assigns a weight to each codon based on wobble-base codon–anticodon interaction rules, comparing it to all synonymous codons. This measure estimates the adaptability of a viral gene to the host tRNA pool ^52^. The tAI weight is calculated as follows:

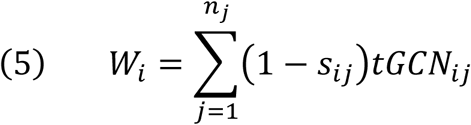

where tRNA gene copy number (tGCN) is the estimated abundance of the *j-*th tRNA molecule recognizing the *i*-th codon, *s*_*ij*_ is a constraint reflecting the affinity between the codon *i* and anticodon *j*, and *n*_*j*_ is the number of anticodons that recognize the *i-*th codon. The tAI weight is normalized by the maximal codon weight:

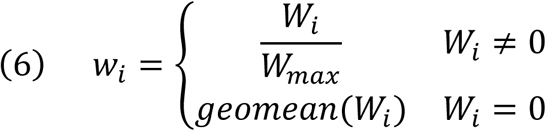

where *W*_*max*_ corresponds to the most optimal codon with the highest weight, among the 61 codons (excluding the stop codons).

#### Supply-to-Demand Adaptation (SDA)

This index measures the balance between codon supply (tRNA abundance) and codon demand (codon usage), allowing consideration of both factors in the translation process. The index is calculated as follows:

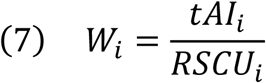

where *tAI*_*i*_ is the <u>non-normalized</u> weight for codon *i* that was calculated in Equation 5, and *RSCU_i_* is the RSCU for codon *i* as it was calculated in Equation 2. The weight was normalized relatively to the most frequent synonymous codon:

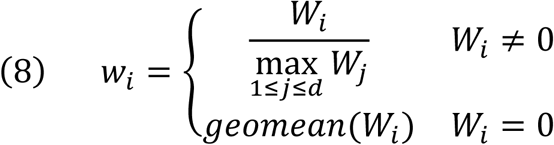

where d is the number of synonymous codons encoding to the same amino acid as codon *i*.

The SDA score for each viral gene is calculated as a geometric mean based on the weight measured on human data, as follows:

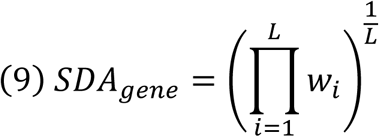

where *L* is the gene length in codons excluding start and stop codons, and *w*_*i*_ are the SDA weights calculated on Equation 8.

### Viral adaptation estimation

To estimate the adaptability of each viral protein to different human tissues, we generated two types of features: Synonymous Dinucleotide Abundance (SDA) and Chimera Average Repetitive Substring (cARS).

SDA scores for each viral protein were computed across 22 tissues (excluding GBM). RSCU values were derived from the tissue-specific codon usage described in Equation 1. Tissue-specific tRNA quantifications were used to compute tAI weights, which were then combined with RSCU values to calculate SDA scores.

#### Chimera Average Repetitive Substring (cARS)

The ChimeraARS algorithm measures the mean length of the longest common substring between a given sequence and a reference. For each position in the sequence, the algorithm identifies the longest matching substring with the reference genome, and the average length across all positions ^43^. The cARS score is defined as:

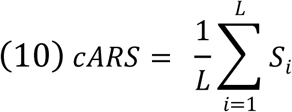

when L is the sequence length and *S*_*i*_ is the length of the longest substring starting at position i.

cARS scores for each viral protein were computed using the top 1%, 5%, and 10% most highly expressed human genes in each tissue (based on TCGA expression data) as reference sets. This procedure was repeated for all 23 tissues, producing tissue-specific cARS scores.

### Per-tropism classifiers

SDA and cARS scores were computed for each viral protein across tissues. These features were then used to train classifiers predicting viral protein tropism. For viral proteins with defined tropism, we applied the Extreme Gradient Boosting (XGBoost) algorithm ^53^.

Two classification strategies were implemented:

1. All-tissues classifier - a single model predicting viral protein tropism across all six tissue categories simultaneously.
2. One-versus-all classifiers - six separate models, each distinguishing one tropism category from all others.

To address class imbalance, we used stratified 10-fold cross-validation ^54^. Performance of the all-tissues classifier was evaluated using average accuracy across 10 folds. To account for class imbalance, average F1 scores were also computed. Performance of the one-versus-all classifiers was evaluated using average accuracy across the 10 folds. In addition, average recall was computed to assess the rate of false negatives and the ability of the models to detect positive cases. Each classifier was trained twice: once using only SDA features and once using both SDA and cARS features.

## Results

### Viruses are adapted to their hosts regulatory mechanisms in a target tissue

We calculated tissue-specific SDA and cARS scores for each viral protein in our dataset and trained two XGBoost classifiers to predict tissue tropism across all tissues: one using only SDA and another incorporating both SDA and cARS. The cARS were calculated using the top 1% of the most highly expressed human genes as a reference. Using 10-fold cross-validation, the SDA-only model achieved an accuracy of 0.786 ± 0.016 and an F1 score of 0.767 ± 0.017. Adding cARS improved performance to an accuracy of 0.817 ± 0.015 and an F1 score of 0.804 ± 0.015. These results indicate that cARS provides additional predictive information relevant to viral protein adaptation. In addition, the lower F1 relative to accuracy reflects class imbalance in the dataset.

Furthermore, the confusion matrix for the SDA+cARS classifier (Figure 1) shows that most tissues are correctly classified, with fewer misclassifications compared to the SDA-only model (Figure S1). It also demonstrates uneven representation across classes, supporting the interpretation of lower F1 relative to accuracy.

**Figure 1.**
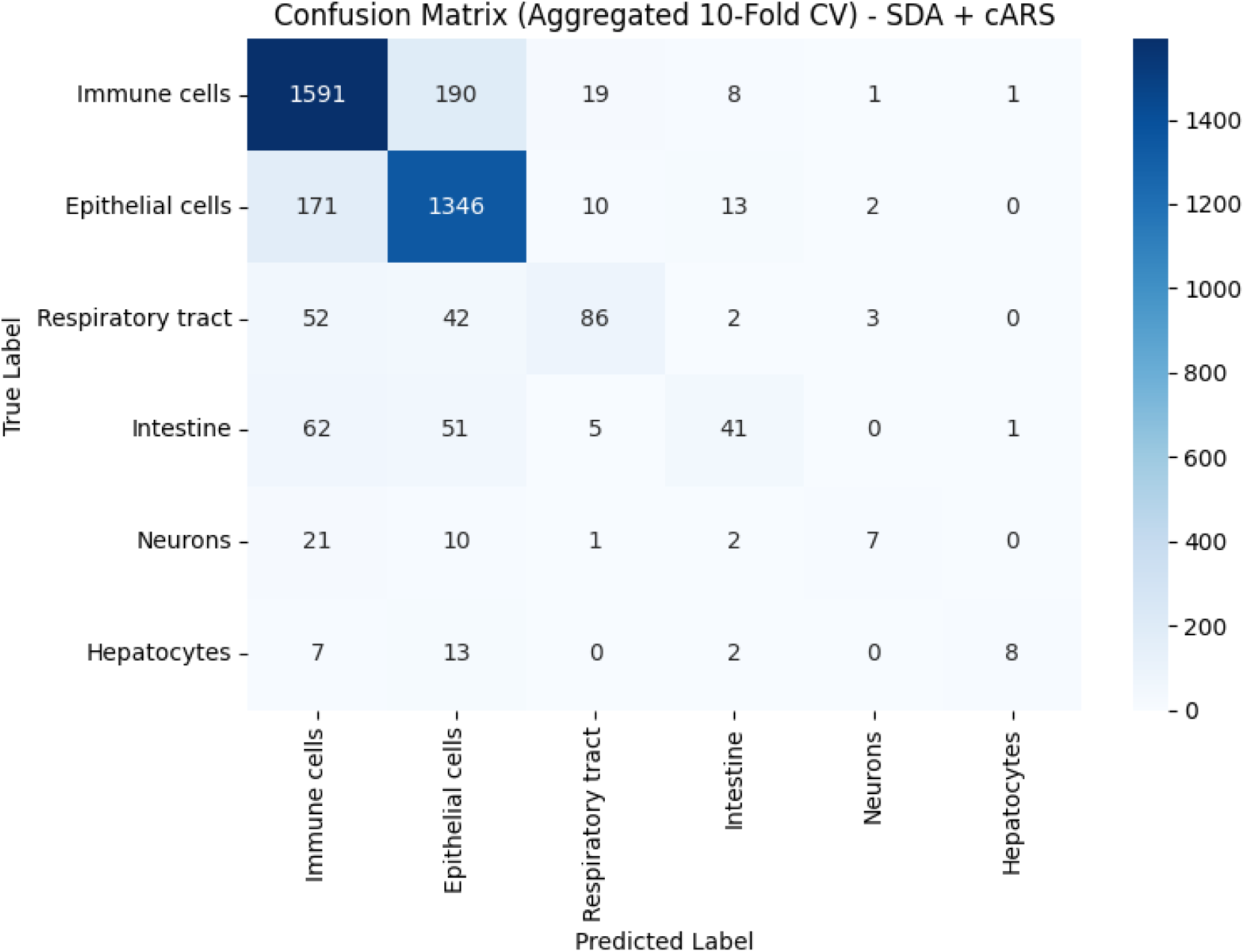

To further examine the contribution of cARS at the tissue level, we next trained one-versus-all classifiers for each tissue and compared the performance of models using only SDA with those incorporating both SDA and cARS. Similar to the all-tissues model, adding cARS consistently improved predictive performance across most tissues. The highest accuracy was for epithelial cells (0.936 ± 0.003), followed by immune cells (0.934 ± 0.002), respiratory tracts (0.910 ± 0.014), hepatocytes (0.899 ± 0.057), neurons (0.874 ± 0.051) and intestine (0.838 ± 0.022), as shown in Figure 2. Compared to SDA only, the model which includes SDA+cARS performed better for all tissues (Table 1).

**Figure 2.**
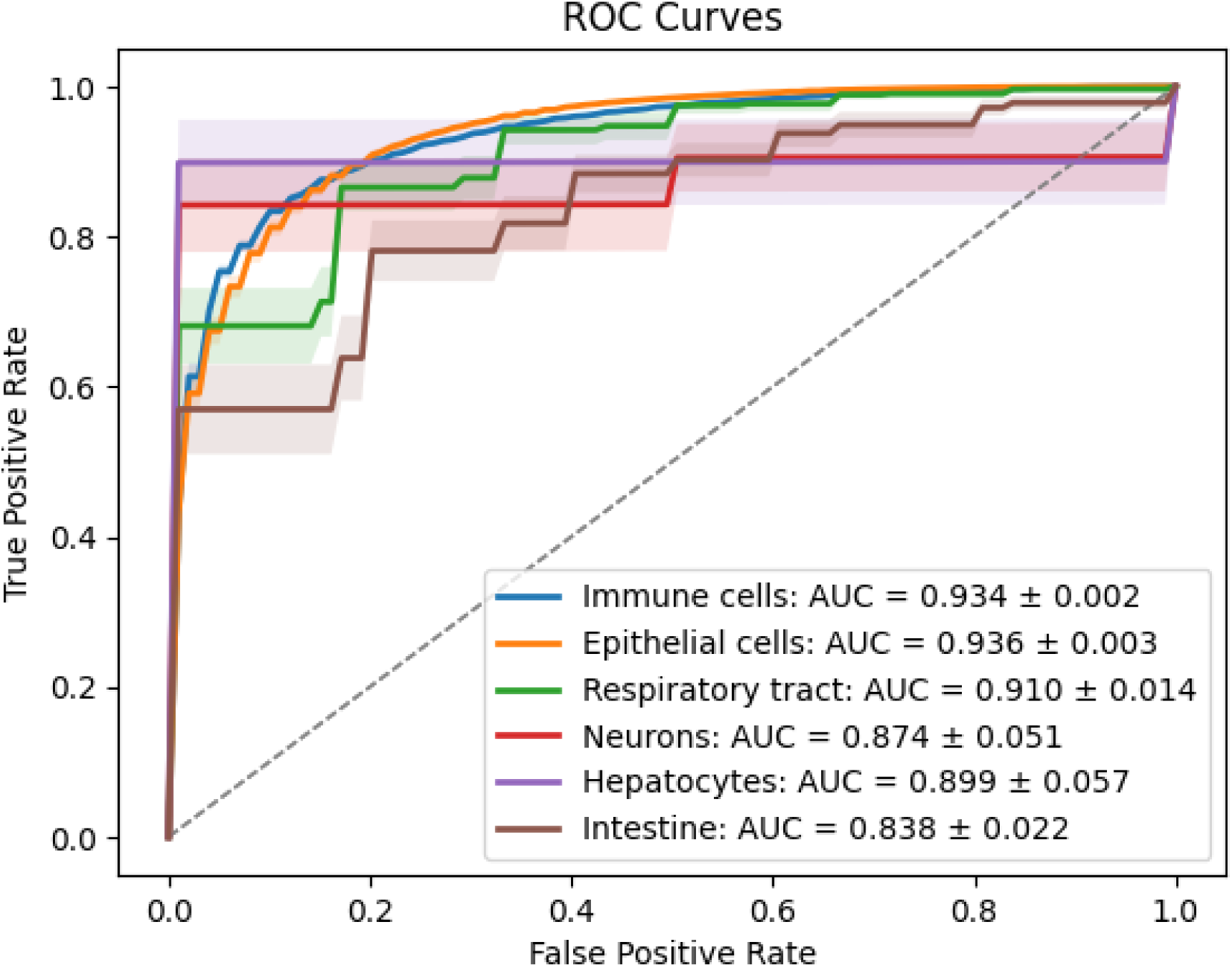

**Table 1.**

|  | SDA only | SDA+cARS |
| --- | --- | --- |
| Epithelial Cells | $0.922 \pm 0.003$ | $0.936 \pm 0.003$ |
| Hepatocytes | $0.871 \pm 0.056$ | $0.899 \pm 0.057$ |
| Immune cells | $0.904 \pm 0.002$ | $0.934 \pm 0.002$ |
| Intestine | $0.782 \pm 0.027$ | $0.838 \pm 0.022$ |
| Neurons | $0.855 \pm 0.055$ | $0.874 \pm 0.051$ |
| Respiratory tracts | $0.842 \pm 0.021$ | $0.910 \pm 0.014$ |

The models (all tissues and one-versus-all) were also trained for cARS with top 5% and top 10% of highly expressed human genes as a reference. All-tissues models have shown an improved performance with less misclassifications (Figures S2-S3), with accuracy of 0.810 ± 0.017 and F1-score of 0.796 ± 0.019 with top 5% as a reference, and an accuracy of 0.815 ± 0.019 and F1-score of 0.802 ± 0.021 with top 10% as a reference.

With one-versus-all model, when using both top 5% as the reference and when using top 10% as the reference improved the results accuracy (Table S1, Figures S4-S5).

## Supplementary Results

**Figure S1.**
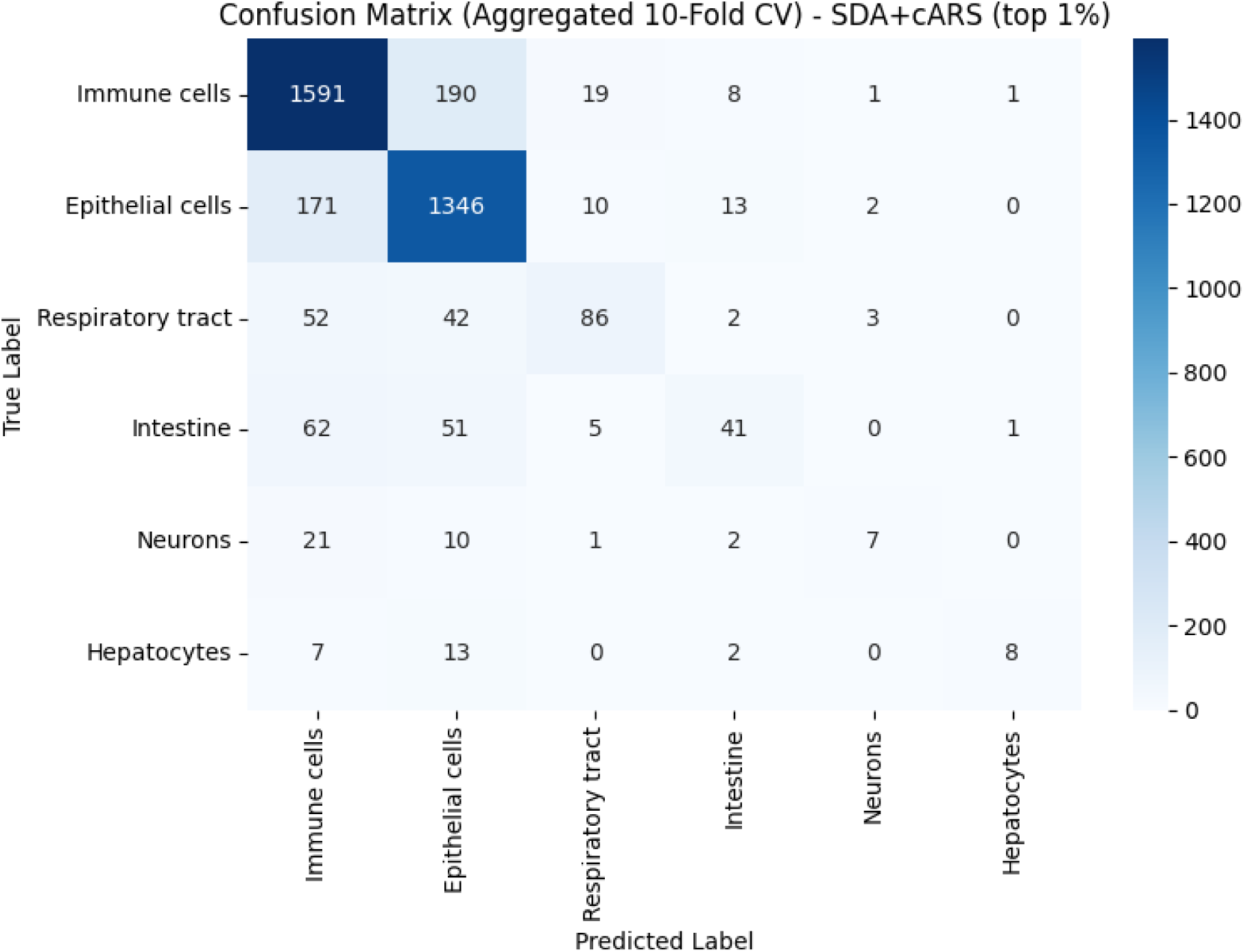

**Figure S2.**
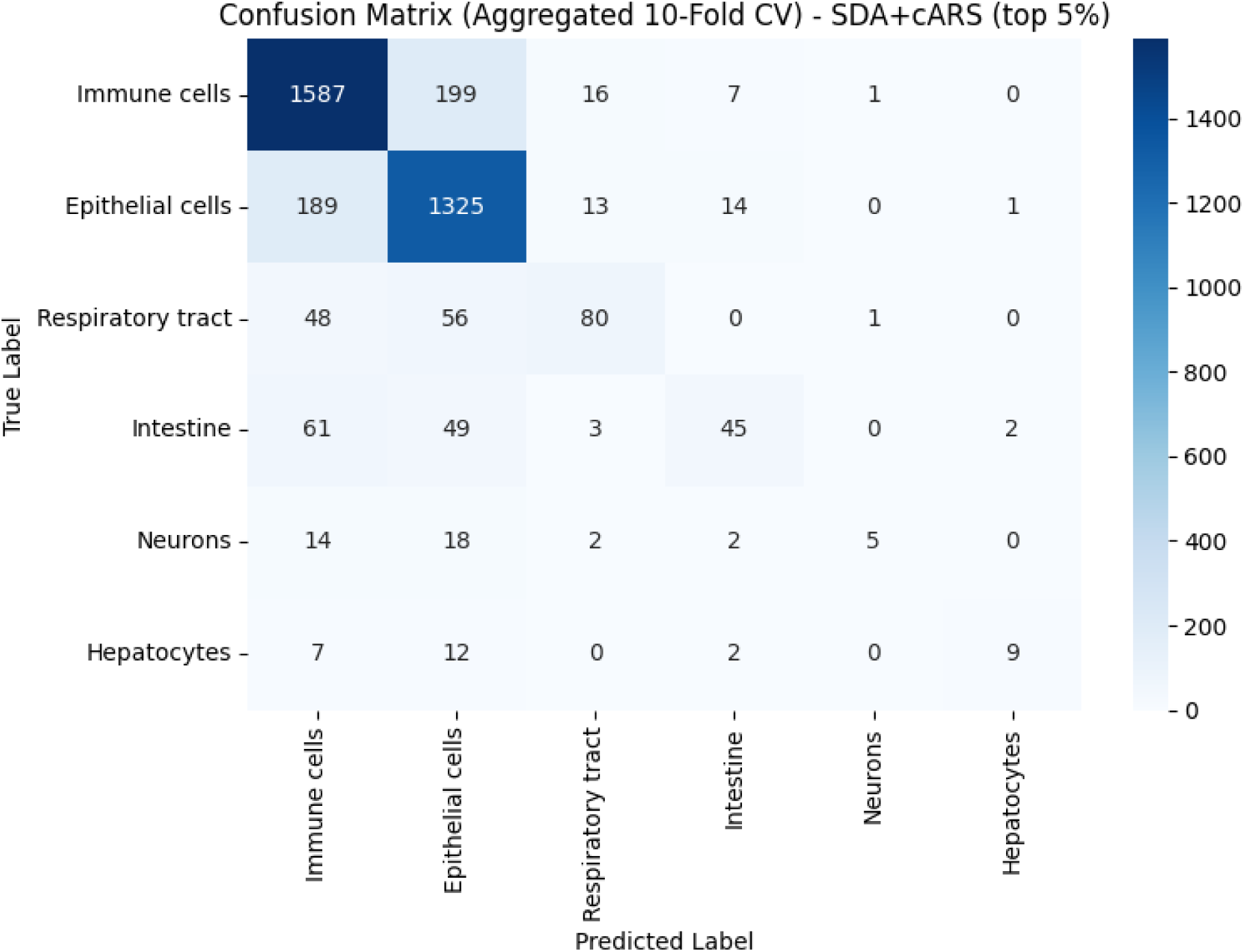

**Figure S3.**
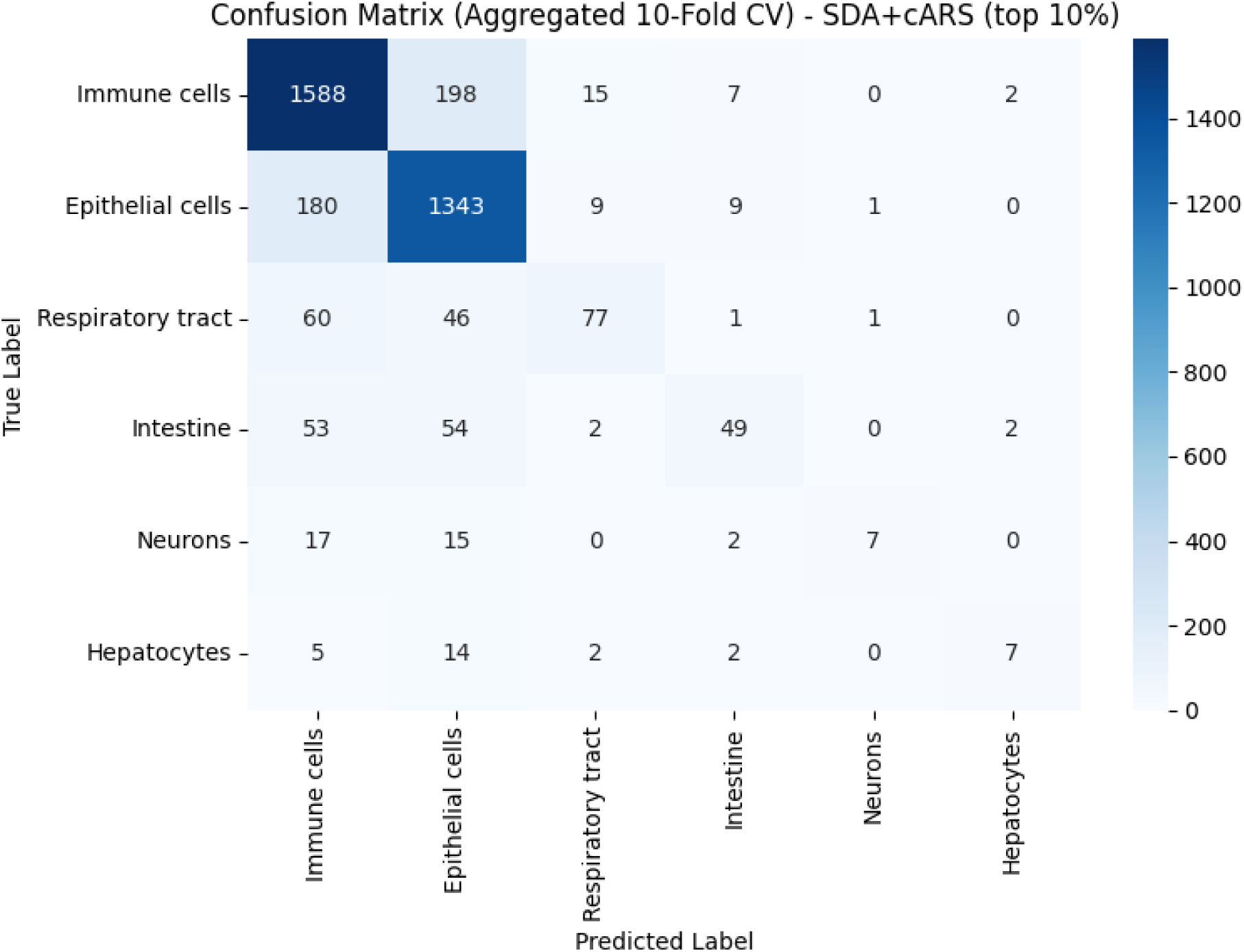

**Figure S4.**
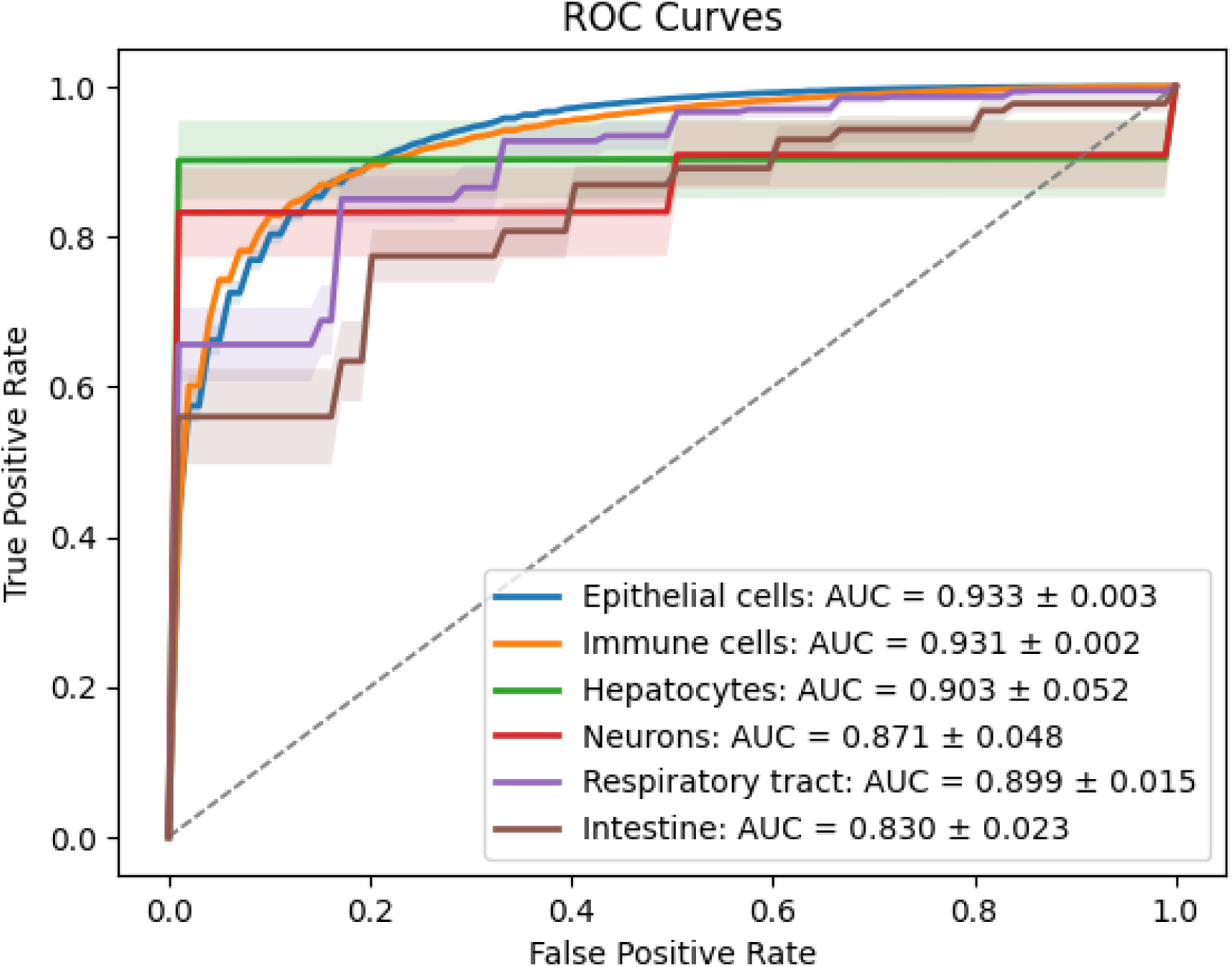
(top 5%)

**Figure S5.**
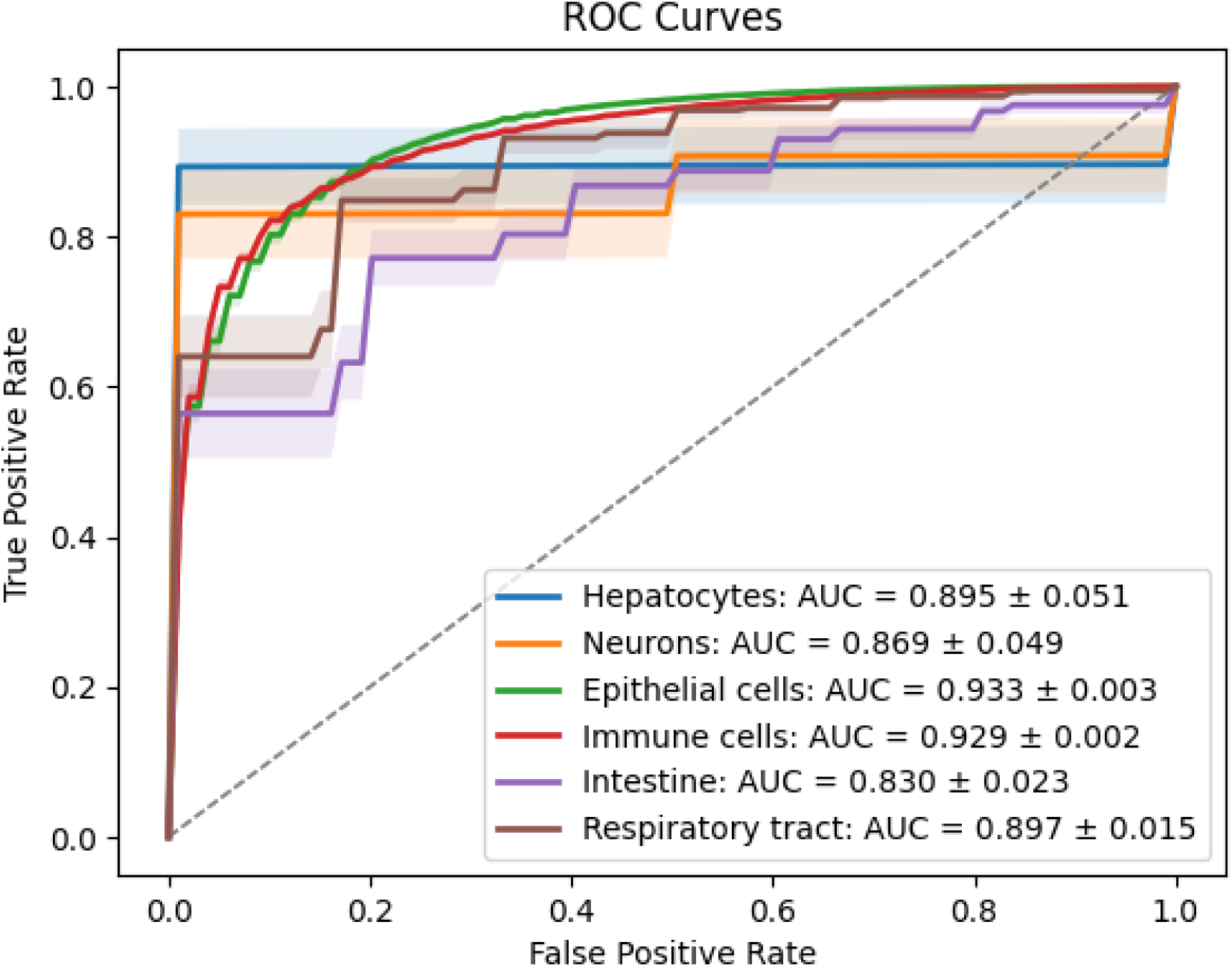
(top 10%)

**Table S1.**

|  | SDA only | SDA+cARS (top 1%) | SDA+cARS (top 5%) | SDA+cARS (top 10%) |
| --- | --- | --- | --- | --- |
| Epithelial Cells | $0.922 \pm 0.003$ | $0.937 \pm 0.002$ | $0.933 \pm 0.002$ | $0.932 \pm 0.003$ |
| Hepatocytes | $0.871 \pm 0.056$ | $0.901 \pm 0.056$ | $0.905 \pm 0.059$ | $0.892 \pm 0.054$ |
| Immune cells | $0.904 \pm 0.002$ | $0.934 \pm 0.002$ | $0.931 \pm 0.002$ | $0.929 \pm 0.002$ |
| Intestine | $0.782 \pm 0.027$ | $0.839 \pm 0.020$ | $0.830 \pm 0.025$ | $0.833 \pm 0.021$ |
| Neurons | $0.855 \pm 0.055$ | $0.872 \pm 0.053$ | $0.870 \pm 0.050$ | $0.866 \pm 0.049$ |
| Respiratory tracts | $0.842 \pm 0.021$ | $0.905 \pm 0.015$ | $0.898 \pm 0.014$ | $0.897 \pm 0.014$ |

